# Mapping the ligand binding landscape

**DOI:** 10.1101/346817

**Authors:** Alex Dickson

## Abstract

The interaction between a ligand and a protein involves a multitude of conformational states. To achieve a particular deeply-bound pose the ligand must search across a rough free energy landscape, with many metastable minima. Creating maps of the ligand binding landscape is a great challenge, as binding and release events typically occur on timescales that are beyond the reach of molecular simulation. The WExplore enhanced sampling method is well-suited to build these maps, as it is designed to broadly explore free-energy landscapes, and is capable of simulating ligand release pathways that occur on timescales as long as minutes. WExplore also uses only unbiased trajectory segments, allowing for the construction of Markov state models (MSM) and conformation space networks that combine the results of multiple simulations. Here we use WExplore to study two bromodomain-inhibitor systems using multiple docked starting poses (Brd4-MS436 and Baz2B-ICR7), and synthesize our results using a series of MSMs using time-lagged independent component analysis. Ranking the starting poses by exit rate agrees with the crystal structure pose in both cases. We also predict the most stable pose using the equilibrium populations from the MSM, but find that the prediction is not robust as a function of MSM parameters. The simulated trajectories are synthesized into network models that visualize the entire binding landscape for each system, and we examine transition paths between deeply-bound stable states. We find that, on average, transitions between deeply bound states convert through the unbound state 81% of the time, implying a trial-and-error approach to ligand binding. We conclude with a discussion of the implications of this result for both kinetics-based drug discovery and virtual screening pipelines that incorporate molecular dynamics.

## I. INTRODUCTION

Much effort in computational medicinal chemistry is devoted to finding *the* correct pose for a given ligand. Docking algorithms examine a large set of possible binding modes, and use empirical scoring functions to determine which of these has the lowest free energy. The accuracy of a given docking algorithm is typically tested by measuring the root mean squared distance (RMSD) between the lowest free energy pose to a single crystal structure pose for a large ensemble of protein-ligand systems [1–3]. Such pose predictions can be very important for the development of lead molecules in the drug discovery process, as they provide an intuition for structure-activity relationships. However, in general, multiple binding poses can exist with similar probabilities – especially in non-optimized protein-ligand systems during screening – and a single-pose paradigm can neglect valuable information that can aid the drug discovery process. For instance, alternative poses with slightly higher free energies can be stabilized during ligand design [4]. Also, the connectivity between states can be taken into account to design inhibitors with long residence times [5, 6], increasingly seen as a desirable objective in drug design [7, 8].

The set of all binding poses can be viewed as points on a multi-dimensional ligand binding free-energy landscape that each occur with some finite probability [9]. A map of this landscape that shows a large ensemble of possible poses and how they are connected with both each other and the unbound state would be a valuable tool in ligand design. A complete, accurate map would allow for the prediction of binding mechanism, free energy and kinetics, and could be used to predict how the stability of these states would change as chemical modifications are made to the ligand. Unfortunately, construction of these maps is challenging, as structural data from experiment is limited to only the most stable states, and molecular simulation is easily trapped by deep metastable free-energy minima.

Recent progress in both hardware and software for molecular simulation is providing our first glimpses to the paths of ligand (un)binding and their kinetics [10]. Ligand release kinetics can be efficiently predicted in some cases using random acceleration molecular dynamics [11], however this method cannot produce absolute binding kinetics, and the external force can affect the ensemble of pathways in ways that are hard to predict. Ligand release paths have been determined using the metadynamics method, where a biasing potential is introduced along an order parameter that describes the binding or release process [12]. This has allowed for the characterization of extremely rare ligand release events, such as the unbinding of dasatinib from Src kinase with a mean first passage time (MFPT) of ≈ 20 s [13], and the unbinding of a Type II inhibitor from p38 MAP kinase, with a MFPT of ≈ 7 s [14]. Although techniques have been developed to subtract the effect of biasing forces on estimates of free energy and kinetics [15], there is no clear way to subtract the impact of the biasing force on the transitions between different microstates.

A number of enhanced sampling methods for ligand-protein interactions employ only unbiased dynamics, which are suitable for building maps of the binding landscape. The adaptive multilevel splitting method uses a series of loops that begin and end in the bound state, progressing farther and farther towards the unbound state as the simulation progresses [16]. This has been used to study the release of benzamidine from trypsin, which has a MFPT of 1.6 ms [17]. Adaptive Markov state modeling, in which on-the-fly Markov state models are used to direct the seeding of new simulations, has also been used on the trypsin-benzamidine system [18]. Another method, weighted ensemble (WE) [19], uses a set of parallel trajectories is balanced between regions of space using cloning and merging operations. This has been used in conjunction with the Northrup-Allison-McCammon method [20] to estimate binding rates using Brownian dynamics [21] and coarse-grained models [22].

The WExplore enhanced sampling method [23] is a variant of WE that has been used to study long timescale ligand release processes. WExplore is particularly suited to study high-dimensional systems, as it builds a set of hierarchical Voronoi polyhedra on-the-fly to divide the space into regions, and to guide the cloning and merging operations in the weighted ensemble framework. WExplore has characterized unbinding paths of a series of molecular fragments from the FK-506 binding protein [24], and has extensively sampled the trypsin-benzamidine system, discovering three distinct ligand release pathways [25]. This method was also used to sample both binding and release pathways of a series of host-guest systems for the SAMPL6 challenge [26], with release pathway timescales as long as 830 s. Finally, Wexplore studied release pathways of the TPPU ligand from the enzyme soluble epoxide hydrolase, experimentally determined to have a MFPT of 660 s. WExplore is thus unique in that it is both 1) built on unbiased dynamics, and 2) capable of sampling ligand release events occurring on timescales of seconds to minutes.

Here we use WExplore to sample the ligand binding landscapes for two protein-ligand complexes, and test its ability to predict global free energy minima when starting from inaccurate starting points. We focus on two bromodomains: small epigenetic reader domains composed of four left-handed alpha helices, which recognize *ϵ*-N-acetylated lysine residues. There are currently 61 known distinct human bromodomains that are contained in 46 different proteins [27]. Bromodomains have been of intense therapeutic interest in recent years [28-39], as bromodomain inhibitors have been proposed as treatments for cancer, diabetes and inflammation. Much attention has been given to the bromodomain and extraterminal (BET) family, including Brd4 for which the first inhibitor was discovered in 2010 [40].

The first complex studied here involves Baz2B (bromodomain adjacent to zinc finger domain protein 2B), which has a relatively small lysine binding pocket, and is considered one of the least-druggable bromodomain targets [28]. Drouin et al [33] developed a chemical probe, “ICR”, that is selective for bromodomains BAZ2A and BAZ2B. We study here the interaction between BAZ2B and an intermediate compound “ICR7” (PDB: 4XUB), which has slight differences from ICR, and is approximately 10-fold less potent for BAZ2B binding (IC50 ≈ 1.1 *µ*M). We also examine the well-studied BRD4 protein, which has two distinct bromodomain subunits. Zhang et al [29] developed an inhibitor “MS436” that preferentially binds to the first bromodomain (Brd4(1)) over the second (*K_i_* ≈ 40 nM). The interaction of MS436 with Brd4(1) (PDB: 4NUD) is the second complex studied here.

Starting from two very different docked poses for each system, we use WExplore molecular dynamics sampling to generate unbinding pathways. We obtain pathways that connect the docked bound states to alternative binding poses, as well as quasi-bound and unbound states. The simulation results are then synthesized into Markov state models that are visualized as networks, and used to predict the globally stable bound poses, which can differ from both starting points. Transition paths that connect the two ligand orientations are analyzed, and we quantify the fraction of transition paths that interconvert on the protein surface, versus interconverting in the bulk. We conclude with a discussion of the implication of these results for the screening of future bromodomain inhibitors.

## II. METHODS

### A. Docking

Docking is performed using Autodock Vina [3] and AutoDockTools from MGLTools package 1.5.6. Coordinates of the protein are taken from Protein Data Bank files 4XUB [33] and 4NUD [29], for Baz2B and Brd4, respectively. We use a cubic grid with 80^3^ points, 0.375 Å spacing, and a center taken from the center of geometry of the ligand in the crystal structure. Crystallographic waters are not included in the docking procedure. We retain the top nine models and use these to choose two starting structures for each protein-ligand system that we label poses A and B. More on how these poses are chosen is given in Section III A.

### B. Molecular dynamics sampling

The CHARMM36 forcefield is used for all minimization and molecular dynamics simulation. Ligands are parameterized with CGenFF [41, 42] and four systems total are built for proteins Baz2B and Brd4, each with poses A and B. Each system is solvated with a cutoff of 12 Å, and ions are added to neutralize the system: for Brd4, three chlorine atoms are added, and for Baz2B, no ions are required. The systems are then energy minimized using harmonic restraints on the protein and ligand atoms: 500 steps of steepest-descent followed by 500 steps of the adopted basis Newton Raphson method. The minimization is then repeated with the restraints removed. Each system is heated gradually from 50 to 300 K by increments of 25 K, with 5000 dynamics steps at each temperature. The systems are then equilibrated with 500 ps of simulation, and the resulting conformation is used to initialize our WExplore simulations. We use a 2 fs timestep for all simulations performed here.

### C. WExplore

WExplore [23] is an enhanced sampling technique built on the weighted ensemble method [19]. In this technique, an ensemble of trajectories are run forward in time and are periodically managed by a central process that can clone or merge trajectories in order to sample over a set of regions as evenly as possible. A set of regions are defined in conformation space, and trajectories are cloned in underrepresented regions (e.g. saddle points), and merged in overrepresented regions (e.g. high probability basins of attraction). The weighted ensemble method has largely been implemented by defining these regions along one or two order parameters [22, 43–45], and the key advance of the WExplore method was to define regions in a high-dimensional order parameter space using hierarchical Voronoi polyhedra (for more information, see [23]). We have found WExplore to be useful for discovering new regions of conformational space [46], and it works best for low-entropy to high-entropy transitions, such as ligand unbinding pathways [10, 24, 25, 47].

In WExplore, trajectories are assigned to regions using distance measurements to a set of characteristic conformations of the system (called “images”), which are dynamically defined over the course of the simulation. To measure distances between conformations we first align the two conformations using a set of residues in the binding pocket that are within 8 Å of the ligand in its initial pose, the distance between the conformations is then the RMSD between the sets of ligand atoms, without any further alignment. This captures ligand rotation, translation and reorganization, and is suitable for building diverse ensembles of ligand-bound poses. As in previous work, we use a four-level region hierarchy, with critical distances of 2.5, 3.5, 5.0 and 10 Å [25, 47], we also use 10000 dynamics steps (Δ*t* = 20 ps) between cloning and merging operations.

This method bears some similarity to Markov state modeling approaches as both are run using entirely unbiased dynamics. However, an advantage of the weighted ensemble family of methods is that no Markovian assumption is needed in order to estimate observables, such as the unbinding trajectory flux that determines the un-binding rate, *k*_off_. This made possible by assigning a statistical weight to each trajectory that governs how strongly it contributes to statistical averages. When a trajectory is cloned, its weight is divided among the clones, thus conserving probability. When two trajectories A and B are merged, the resulting trajectory C has weight *w_C_* = *w_A_* + *w_B_*, and it takes on conformation A with probability *w_A_/w_C_*, and conformation B with probability *w_B_ /w_C_*. Here, following previous work [25], to improve sampling efficiency we impose maximum and minimum weights that walkers can achieve: *w*_max_ = 0.1 and *w*_min_ = 1*e*^−12^. This is enforced by disallowing cloning or merging operations that would violate these rules. Here we use both the statistical weights of the walkers to calculate exit rates, as well as weights from Markov models to predict globally stable bound poses.

For each of the four systems we perform three WExplore simulations, with 48 trajectories each. These are run for 730 cycles, which are comprised of 20 ps of sampling followed by merging and cloning operations. In aggregate, we report on 701 ns per WExplore simulation, or 8.4 *µ*s total.

### D. Exit rates

To calculate the exit rates we employ an ensemble splitting strategy [48], where two basins (bound and unbound) are used to define binding and unbinding ensembles. Our simulations are run in the unbinding ensemble, where trajectories are initialized in the bound basin and are terminated in the unbound basin. The trajectory flux from the unbinding ensemble into the binding ensemble is equal to the unbinding rate (*k*_off_), and can be calculated as the sum of the weights of the exiting trajectories divided by the elapsed time. We have found previously that although these measurements can vary significantly between WExplore simulations, the average over an ensemble of simulations can compare well to experimentally determined off-rates [25, 47].

### E. Markov state model and network modeling

As our simulation results broadly sample the ensemble of ligand bound poses, including complete exit trajectories, there is considerable overlap between those that start in Pose A and those that start in Pose B. We use co-clustering to synthesize the two datasets, and use these models to visualize our pose network, predict globally stable states, and determine properties of A ↔ B transition ensembles.

For the pose networks we cluster using a set of ligand-protein distances. This allows all poses to be distinguished on a structural basis, creating a detailed view of the free energy landscape. For both Baz2B and Brd4, the set of distances was constructed using two sets of atoms (a set of selected ligand atoms and the set of protein C*α* atoms that are within 20 Å of the center of mass of the ligand), and choosing every possible combination of atoms between the two sets. The selected atoms for each ligand was a set of heavy atoms, 13 for MS435 and 15 for ICR7, selected manually, that cover all functional groups of the ligand (Figure S1). The set of ligand-protein distances (962 for Brd4-MS435 and 1035 for Baz2B-ICR7) is used as a base set of features to describe our dataset.

We then use time-lagged independent component analysis (tICA) [49, 50] to identify coordinates that change slowly as a function of time. As WExplore uses a set of trajectories that are cloned and merged at each time step, knowledge of the trajectory history is required for the tICA analysis. This was accomplished by construction of a branching tree of trajectories using the wepy package [51], allowing each point to be traced backward in time in a unique fashion. Many tICA clustering parameters were used in this work to examine the robustness of our cluster predictions. We varied the number of dimensions used in tICA analysis (*n_tICA_* = 3, 5, 10), the tICA lag-time (*τ_tICA_* = 0.2, 1 ns), the Markov model lag-time (Δ*t* < *τ* < 100Δ*t*), and the number of clusters (*n_c_* = 500, 800, 1200). As the initial sampling is heavily concentrated around the initial points chosen from docking, we also discard a fraction of the initial data (0 *< f_d_ <* 0.8). Parameters used in specific tICA cluster sets are given in Table I, and for each set we construct 63 Markov models (7 *τ* values and 9 discard fractions). We construct another 63 Markov models without tICA clustering for each of *n_c_* = 500, 800 and 1200. These use k-means clustering directly on the set of ligand-protein distances. In total, we compute predictions of the globally stable state using 945 different Markov models for each of the Brd4 and Baz2B systems.

**TABLE I.** Parameter sets used for clustering with tICA.

| $n_{tICA}$ | $\tau_{tICA}$ (ns) | $n_c$ |
| --- | --- | --- |
| 3 | 0.2 | 500 |
|  |  | 800 |
|  |  | 1200 |
| 5 | 0.2 | 500 |
|  |  | 800 |
|  |  | 1200 |
| 5 | 1.0 | 500 |
|  |  | 800 |
|  |  | 1200 |
| 10 | 0.2 | 500 |
|  |  | 800 |
|  |  | 1200 |

For the network models, each cluster is represented as a node, and non-zero off-diagonal elements of the transition matrix are represented as edges. As in previous work [25, 47, 52], the weight of an edge between nodes *i* and *j* (*e_ij_*) is as follows:
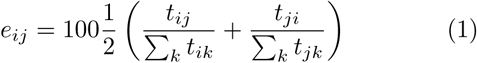

that is, the average of the conditional transition probabilities in either direction between *i* and *j*, multiplied by 100. Determination of transition matrices, weight calculations, and graph construction were performed with the CSNAnalysis package v0.5 [53]. Network layouts are created in Gephi using the Force Atlas algorithm, with repulsion strength 200, attraction strength 10 and maximum displacement 10. The layout is minimized first while allowing for node overlap, followed by brief a minimization while preventing overlap, with the maximum displacement reduced to 1.0.

## III. RESULTS

### A. Starting poses

We use Autodock Vina [3] to generate a set of nine possible bound poses for both Baz2B and Brd4 (Figures S2 and S3). From this set we select two poses that show large differences in their global orientation while maintaining high predicted affinities. These are used as starting points for WExplore MD sampling (Figure 1). RMSD to the crystal structure for all four starting structures ranges from 3.1 Å to 10.5 Å. Pose A for Baz2B has roughly the same orientation as the crystal structure, but is shifted further into the binding pocket. The N6 atom of the methyl-pyrazole group, which in the crystal structure forms a hydrogen bond with ASN1944, is roughly 3 Å deeper, forming a hydrogen bond with the TYR1901 sidechain. The N5 atom on the central imidazole ring moves down to form a similar interaction with ASN1944. Pose A for Brd4 is also shifted approximately 3 Å deeper into the pocket. Pose B in both cases is bound in a completely different orientation. In Baz2B the nitrile group is bound to the recognition pocket, with the triazole and imidazole rings spreading in different directions. In Brd4 the ligand is rotated approximately 180 degrees, with the 2-aminopyridine ring deeply bound in the pocket.

**FIG. 1.**
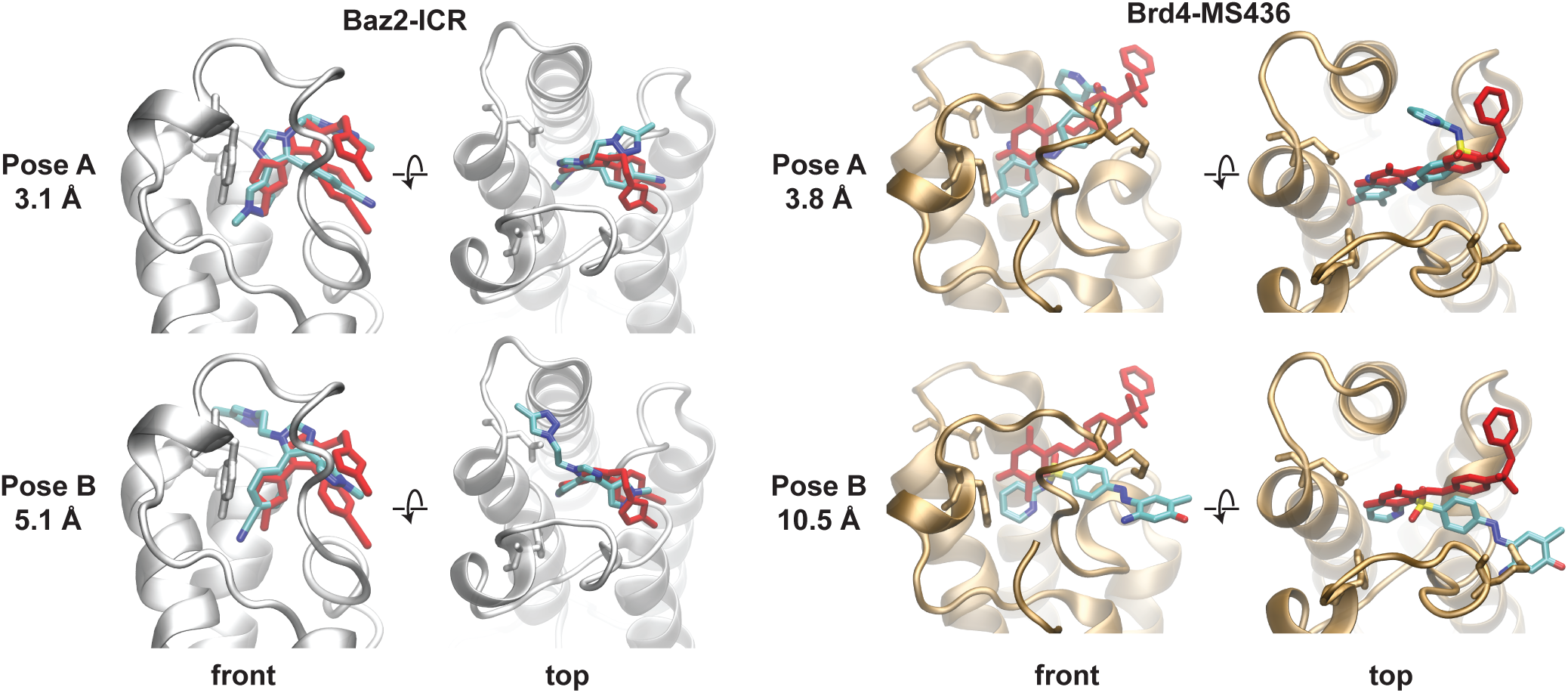
Initial poses from docking. Left: Front and top views of starting poses A and B of the Baz2B-ICR7 system. Residues Tyr1901 and Asn1944 are shown in licorice representation. Right: Front and top views of starting poses A and B of the Brd-MS436 system. Protein residues Tyr97 and Asn140 are shown in licorice, which are homologous to those shown in Baz2B. In both cases, the crystal structure pose is shown in red, and poses A and B are shown with colors according to atom type.

### B. Ranking poses by exit rates

Exit points are obtained in each of the 12 WExplore simulations. We obtain a total of 371 exit points for Baz2B, and 124 for Brd4, which reflects the higher affinity of the MS436 ligand (Table II). These exit points all represent structures that have at least 10 Å of clearance between the ligand and the protein. As seen in Figure 2, they are heterogeneous, and are widely distributed in space surrounding the protein.

**FIG. 2.**
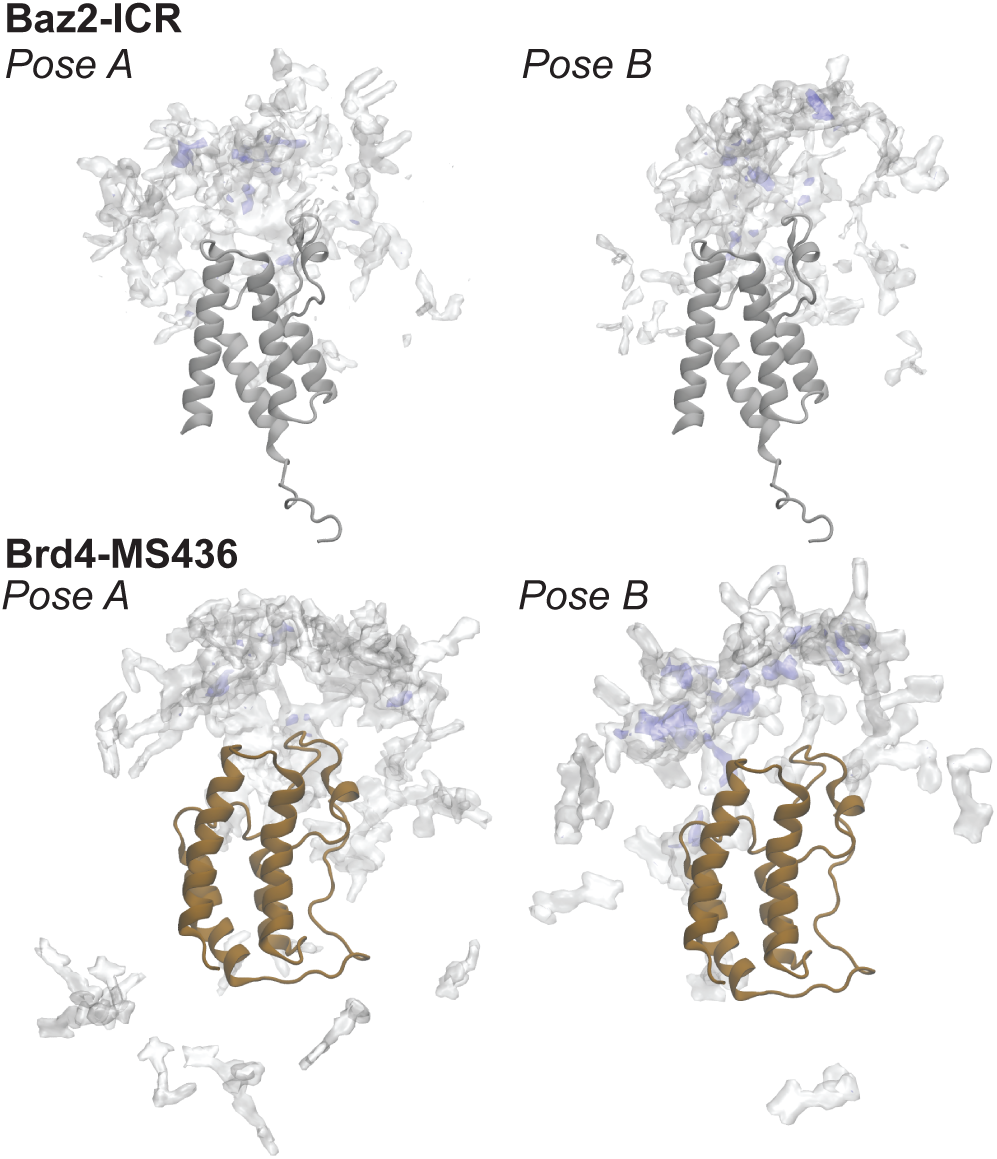
Exit point clouds. For Baz2B-ICR7 (top) and Brd4-MS436 (bottom), we show density distributions for ligand exit points, which are conformations where the closest interatomic distance between the ligand and protein is at least 10 Å. The densities are determined using the VolMap facility in VMD, and are visualized using isovalues 0.02 (transparent white) and 0.08 (transparent blue).

**TABLE II.** Pose-specific dissociation kinetics determined by exit point weights.

| | | Number of exits | | | Total exits | $w_{\text{total}}$ | $k_{\text{off}} \text{ (s}^{-1}\text{)}$ | MFPT<sub>off</sub> |
| --- | --- | --- | --- | --- | --- | --- | --- | --- |
|  |  | Run 1 | Run 2 | Run 3 |  |  |  |  |
| Baz2B | Pose A | 38 | 78 | 53 | 169 | 0.035(0.016) | $2.4(1.1) \times 10^6$ | 420 ns |
| | Pose B | 68 | 58 | 76 | 202 | 0.10(0.04) | $6.6(2.7) \times 10^6$ | 150 ns |
| Brd4 | Pose A | 25 | 28 | 24 | 77 | $5.4(4.2) \times 10^{-8}$ | 3.6(2.9) | 280 ms |
| | Pose B | 24 | 9 | 14 | 47 | $3.2(2.6) \times 10^{-5}$ | $2.2(1.8) \times 10^3$ | 0.46 ms |

Using the sum of the weights of these exit points, we determine off-rates for each protein and starting pose. A small, 3-fold difference was observed in *k*_off_ between ligand poses in the Baz2B system, with Pose A predicted to be slower. For Brd4 we predict Pose A to have a slower off-rate than Pose B by a factor of about 600. Thus, as a pose ranking technique, the exit flux would predict Pose A to be more stable than Pose B in both cases. However, as in previous work, the exit fluxes have high run-to-run variability, as seen by the large error estimates in both *w*_total_ and *k*_off_ in Table II.

It is important to keep in mind that the link between a pose-specific *k*_off_ and the thermodynamic probability of a binding pose is not straight-forward. In other words, long exit times do not imply that a particular pose is stable, or that it is a relevant template to use for drug design. The most probable ligand pose will form a basin of attraction in the free energy landscape, and a set of nearby poses – although they themselves may be unstable – will commit to that basin with high probability, and thus have a similarly long *k*_off_. Thus, to understand the link between pose-specific *k*_off_ and pose probability we must understand how poses are connected to each other, and with the unbound state.

### C. Markov state models for pose prediction

#### 1. Special considerations for Weighted Ensemble datasets

As all of the WExplore simulations are run with the natural energy function (e.g. no biasing forces), we can employ Markov state model analysis to divide our conformation space into a set of states, and estimate their equilibrium probabilities. Although this is possible with weighted ensemble datasets, the directed cloning and merging operations introduce some special considerations into how the transition count matrices should be constructed. To illustrate this, consider a weighted ensemble simulation as a branching “trajectory tree”, with each trajectory growing upward in time (Figure 3). Cloning events can be viewed as branchpoints of this tree, and merging events result in the termination of a branch, with its weight transferred to another point in the tree. For a given transition matrix lag time (*τ* = *n*Δ*t*), a count matrix can be constructed by the set of all possible tree paths *P* = {(*a*_0_ → *b*_0_), (*a*_1_ → *b*_1_)*,…*}, where *a* and *b* are connected by a path of length *n*. So once all of the structural data is clustered, the cluster assignments are used to label the tree, and the transition count matrix *could* be constructed as:
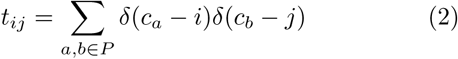

**FIG. 3.**
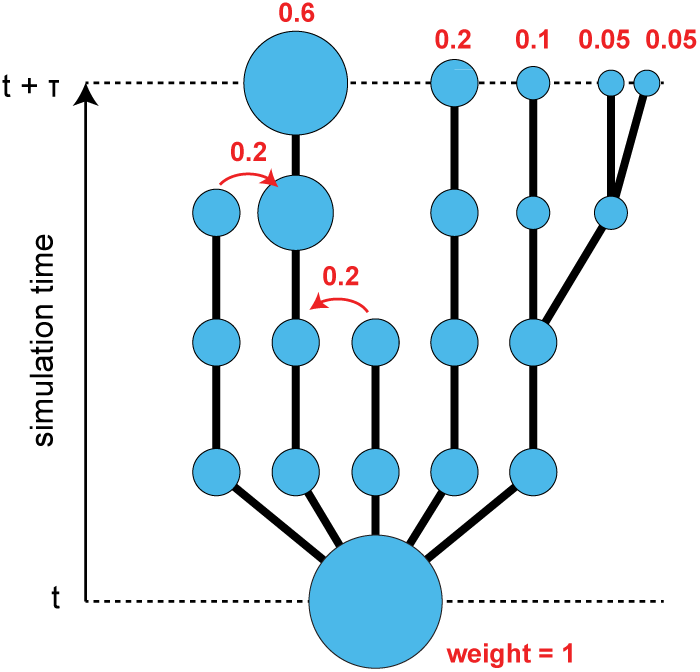
Trajectory tree schematic. Merging and cloning events in weighted ensemble simulations can be represented as a branching tree. Cloning events are branching points, and merging events result in the termination of one branch, and the transfer of its weight to another branch (curved red arrows). To construct a transition count matrix with a lag time of *τ*, all possible complete paths between *t* = *nτ* and *t* = (*n* + 1)*τ* should be considered. There are five such paths in this schematic.

where *c_x_* is the cluster assignment of conformation *x* in the trajectory tree. However, this transition matrix construction is problematic, as it gives all paths equal weight, regardless of the statistical weight of the trajectory determined by the weighted ensemble algorithm. To appreciate this, consider the set of trajectories that begin in the bottom state in Figure 3. The rightmost trajectories have been cloned and would thus be over-represented compared to the others if the transition matrix was constructed according to Eq. 2. In fact, since low-probability unbinding trajectories are systematically amplified in the simulations here, the transition count matrix would be systematically biased toward the unbound ensemble. For this reason it is proper to follow the standard approach of weighted ensemble simulations, where trajectories contribute to observables *according to their statistical weight*.

These weights account for the cloning and merging steps in the WExplore algorithm, and are shown for the final states in Figure 3.
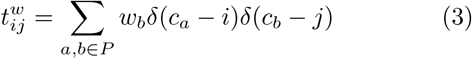

where *w_b_* is the weight of the trajectory at point *b* in the tree. In the next section we compare both of these approaches.

#### 2. Estimates of binding affinity

We construct a series of Markov models for the un-weighted (Eq. 2) and weighted (Eq. 3) transition count matrices. For each model we calculate the equilibrium weights for each state, and estimate the binding affinity using the sum of the probabilities of the unbound states:
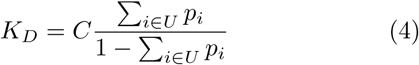

where a state *i* is in *U* if the minimum distance between the ligand and the protein is greater than 5 Å, and *C* is the concentration of ligand, which here is equal to 4.3 mM, calculated as 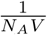, where *N_A_* is Avagadro’s number and *V* is the box volume in liters. The minimum ligand-protein distance for each cluster is determined using an average of 10 randomly chosen structures from each cluster. It is important to note that our simulations are strictly in the unbinding ensemble, that is, trajectories are initialized in the bound state and are terminated once they achieve a minimum ligand-protein distance greater than 10 Å. For this reason, we expect the unbound state to be under-sampled, and we view the *K_D_* calculated by Eq. 4 as a lower bound.

As expected, we observe significant differences between *K_D_* values calculated using the weighted and un-weighted transition count matrices. The *K_D_* values for all parameter sets are shown as heat maps in Figures S4-S7, although they do not vary significantly between the parameter sets. The most significant variation between parameter sets is for the weighted transition count matrices of Brd4-MS436, which has *K_D_* values ranging from 1.3 nM (for *n_tICA_* = 5, *τ_tICA_* = 0.2 ns, *n_c_* = 500) to 820 nM (for *n_tICA_* = 5, *τ_tICA_* = 1 ns, *n_c_* = 500). The average (on a logarithmic scale) minimum and maximum *K_D_* values for the two systems and two count matrices are summarized in Table III. The weighted count matrix shows excellent agreement with the K_i_ for Brd4-MS436. For Baz2B-ICR7, only the IC50 was measured, although the IC50 and K_i_ values were almost equal for a closely related compound [33]. All of the Markov models that we construct predict a binding constant that is higher by at least a factor of 20, which is discussed further in the Discussion section. In both cases the weighted count matrices achieve a much better agreement with experimental binding constants, and these are used for all subsequent analyses.

**TABLE III.**
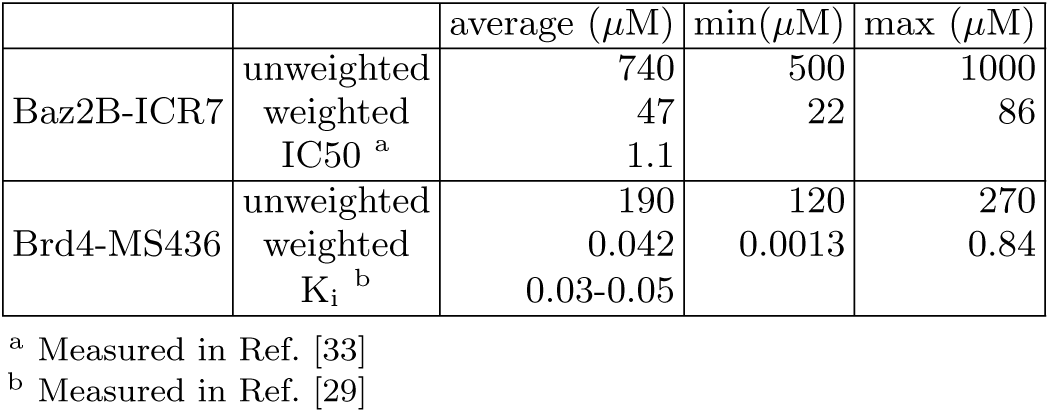
Calculated diffusion constants (*KD*)

| | | average ($\mu\text{M}$) | min($\mu\text{M}$) | max ($\mu\text{M}$) |
| --- | --- | --- | --- | --- |
| Baz2B-ICR7 | unweighted | 740 | 500 | 1000 |
|  | weighted | 47 | 22 | 86 |
|  | IC50 <sup>a</sup> | 1.1 |  |  |
| Brd4-MS436 | unweighted | 190 | 120 | 270 |
|  | weighted | 0.042 | 0.0013 | 0.84 |
|  | K<sub>i</sub> <sup>b</sup> | 0.03-0.05 |  |  |

<sup>a</sup> Measured in Ref. [33]<sup>b</sup> Measured in Ref. [29]

#### 3. Prediction of lowest free energy poses

For each set of Markov model parameters, we lowest free energy pose in each network and calculate the RMSD of this state to the crystal structure. The RMSDs shown are averages over 10 randomly chosen structures for the lowest free energy cluster. These low free-energy pose RMSDs are shown as heat maps for all of the tICA parameter sets examined in Figures S8-S11. RMSD heat maps for two sets of clustering parameters are shown for both systems in Figure 4. Figure 4A shows the results for *n_tICA_* = 5, *τ_tICA_* = 0.2 ns and *n_c_* = 1200. For this set of parameters, structures similar to pose A are not always predicted to be more stable than those similar to pose B, but we find large regions of stability where the lowest free-energy pose has a low RMSD (*<* 2.5 Å) to the crystal structure. In Figure 4B, the number of tICA dimensions is increased from 5 to 10, and the predictions for the most stable pose are changed considerably. In general we find that predictions for the lowest free-energy pose are sensitive to tICA clustering parameters (see Figures S8-S11).

**FIG. 4.**
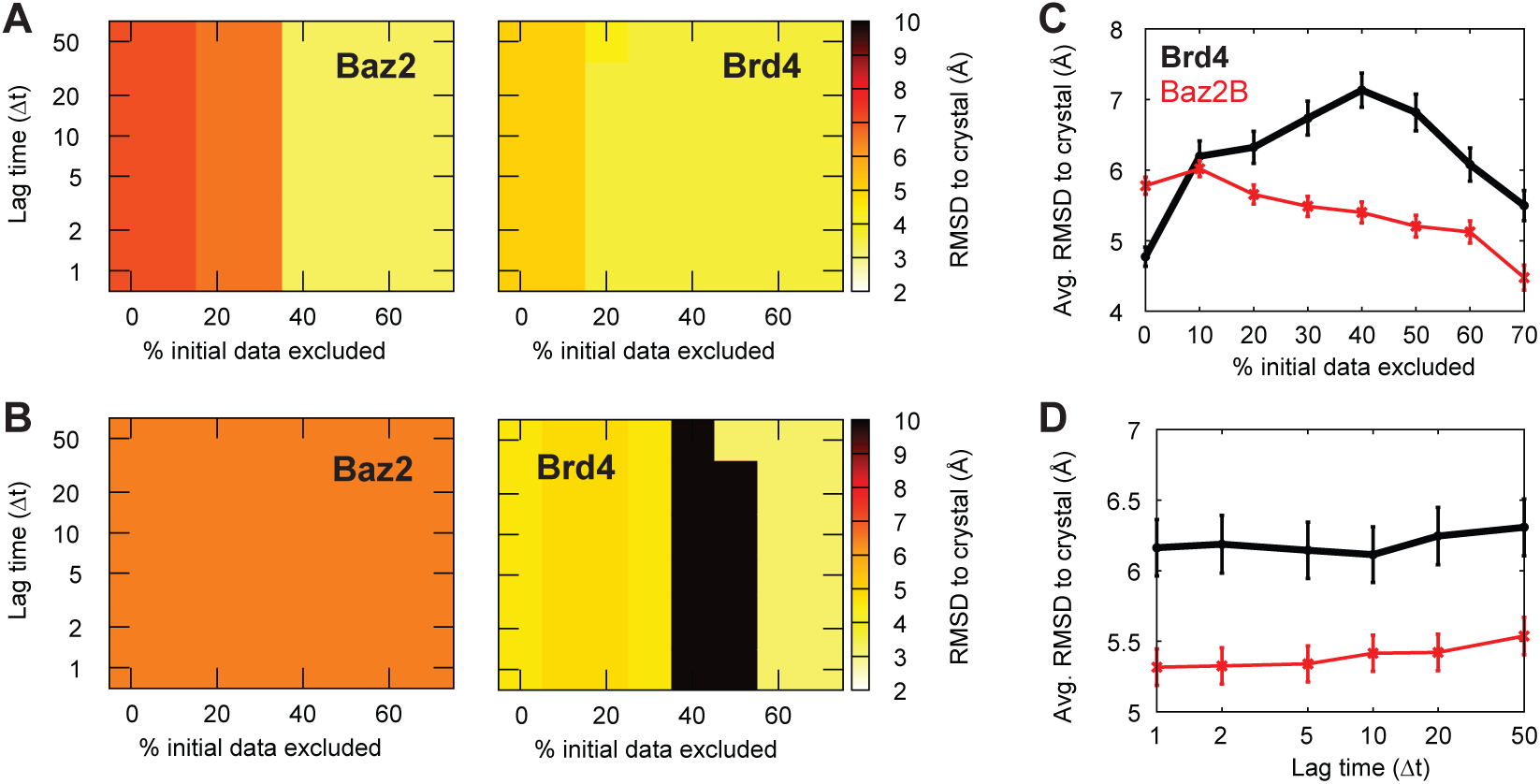
RMSD of lowest free-energy poses. The lowest free energy node is determined for several values of two parameters: the Markov model lag time and the percentage of initial data that is excluded. (**A,B**) Heat maps show the RMSD of the lowest free energy pose determined by the Markov model to the crystal structure. These maps are constructed using *τ_tICA_* = 0.2 ns and *n_c_* = 1200, with *n_tICA_* = 5 for **A** and 10 for **B**. The complete set of maps is shown in Figures S8-S11. The average lowest free-energy pose RMSD is shown as a function of the percentage of initial data discarded (*p_excl_*, **C**), and the Markov model lag time (*τ*, **D**). Points are calculated as the average of all data shown in Figures S8-S11. Error bars show the standard error of the mean. In both panels, Brd4-MS436 data is shown as a thick black line, and Baz2B-ICR7 is shown as a thin red line.

To generally examine whether our pose prediction improves as a function of lag time (*τ*), or the percentage of initial data excluded from analysis (*p_excl_*), we averaged the low free-energy pose RMSDs along these axes. We find a clear trend for Baz2B that increasing predictions improve with increasing *p_excl_* (Figure 4C). For Brd4 we find that the predicted RMSD initially deteriorates and then improves with increasing *p_excl_*. Interestingly, we find that the average quality of predictions for both Baz2B and Brd4 do not improve with longer Markov model lag times (Figure 4D). We also find that we consistently achieve better results for the Baz2B-ICR7 system compared to the Brd4-MS436 system.

To investigate whether the tICA vectors are reporting on long timescale interactions, we use the tICA vectors to color the pose networks described in the next section (Figure S12-S13). In both cases, the first tICA vector (which reports on the longest timescale process) corresponds to transitions between the Pose A region and the rest of the network. The second tICA vector corresponded to transitions between the unbound/quasi-bound states and the deeply bound states. Vectors 3-5 described transitions between disparate communities within the quasi-bound ensemble, the only exception being Baz2B-tICA4, which also describes transitions between two adjacent deeply-bound communities (see Section III E for more details). We conclude that the tICA analysis is working as intended, as the vectors capture the longest timescale motions in our system. To compare, we also generate a separate set of Markov models without using tICA, where the set of ligand-protein distances was directly clustered. This is helpful as it eliminates the need to define *τ_tICA_* and *n_tICA_*. However, we obtain similar mixed results for the RMSD predictions, where only small domains of robustness are found, and different results can be obtained with different numbers of clusters (Figure S14).

To investigate these poses in more detail, we randomly pick a set of 100 structures from the clusters from *n_tICA_* = 5, *τ_tICA_* = 0.2 ns and *n_c_* = 1200 that are predicted to be the most stable for higher *p_excl_*. The median structure from this set (with the lowest ligand RMSD to the other members) is shown along with density maps that are calculated from all set members in Figure 5. We see a significant amount of variation within the set of structures for both systems. For Brd4 we see that the median structure is bound significantly lower than the crystal structure pose, in line with starting pose A. We do not reproduce the orientation of the solvent-exposed pyridine ring observed in the crystal structure, although this could be due to interactions between other copies of Brd4 in the crystal lattice (Figure S15). Poses sampled from the lowest free energy cluster have RMSD values to crystal ranging from 2.4 to 5.5 Å. We find better agreement for the Baz2B-ICR7 system, with RMSD values ranging from 1.2 to 4.9 Å. The median structure reproduces the crystal structure interactions in the binding pocket, and this is consistent across the majority of structures examined. The density becomes more spread out for the solvent-facing triazole ring, which is observed partly in an un-stacked orientation relative to the benzonitrile group. This is in line with NMR measurements of the ICR7 compound, which indicate that the triazole and benzonitrile groups do not exhibit a stacked conformation in solution [33]. However, the lack of stability of the bound, stacked conformation could still indicate that ring stacking interactions are not sufficiently strong in our parameterization of the forcefield for ICR7.

**FIG. 5.**
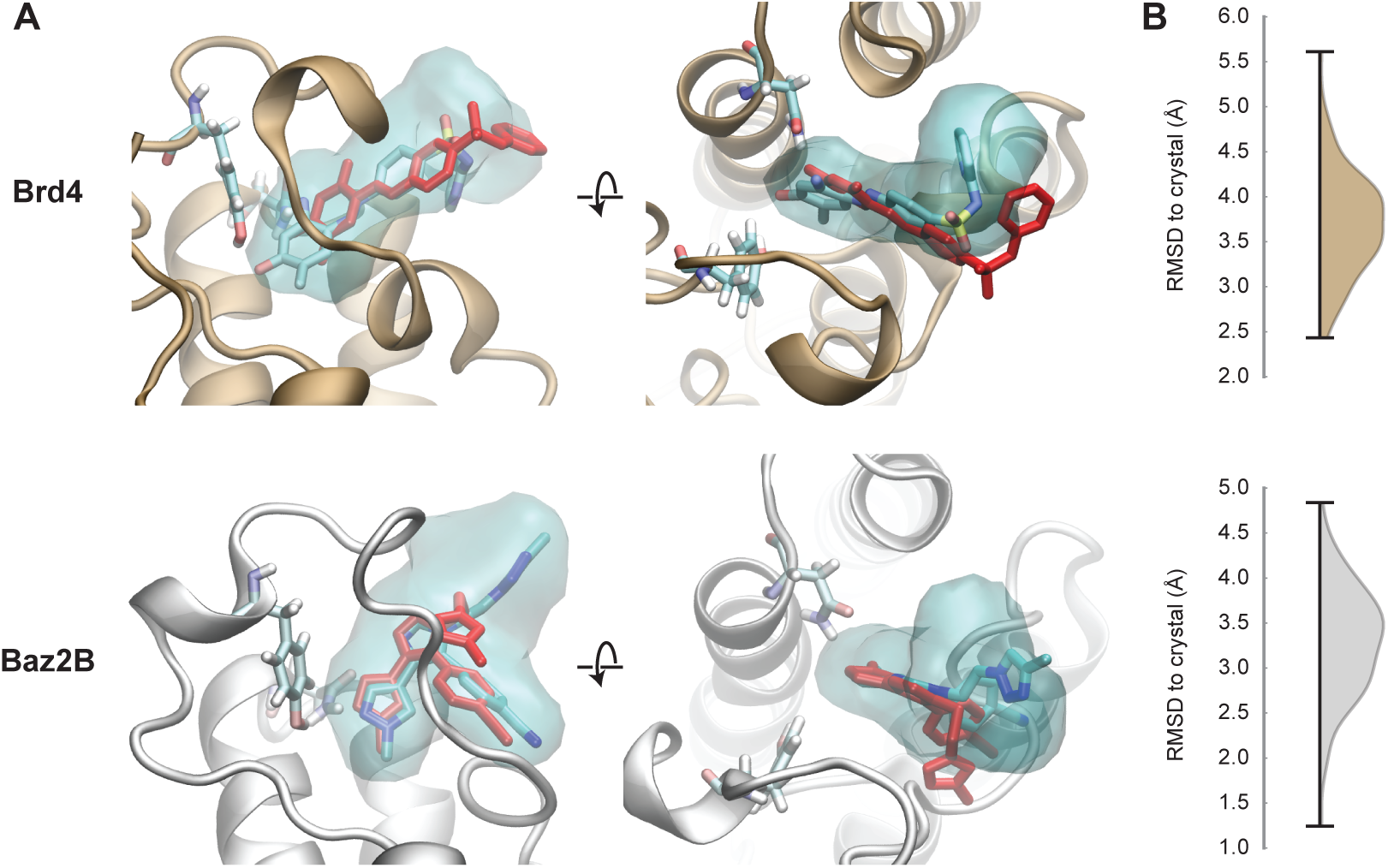
Structures from the lowest free energy clusters. Ligand densities are shown for a set of 100 randomly chosen structures from the nodes identified in Figure 4A. **A** The crystallographic pose is shown in red, and the median ligand pose is shown colored by atom type. Two residues are shown that are important for binding: Tyr97 and Asn140 for Brd4 and Tyr1901 and Asn1944 for Baz2B. The ligand density is determined using the VolMap tool of VMD and is shown at an isovalue of 0.1. **B** The RMSD distribution of the 100 selected poses to the crystal pose is shown for Brd4 (top) and Baz2B (bottom).

### D. Bound pose networks

We use conformation space networks to visualize the landscape of bound poses for both systems (Figure 6). We again focus on a single set of Markov state model parameters (*n_tICA_* = 5, *τ_tICA_* = 0.2 ns, *n_c_* = 1200, *p_excl_* = 70 and *τ* = 0.2 ns), and while the parameters affect the weights of specific states in the network, the overall topology of the network is robust. Edges are given a weight according to Eq. 1, with the weighted transition matrix elements from Eq. 3. For visualization, only edges with a weight greater than 0.4 are shown, and nodes that are not connected to the giant component of the network are discarded. After filtering, the Baz2B network shows 1195 nodes (99%) and 15848 edges (66%), and the Brd4 network shows 1157 nodes (96%) and 7442 edges (49%). These numbers illustrate that the Baz2B system is much more interconnected than Brd4, which is also clear from visual inspection of the networks.

**FIG. 6.**
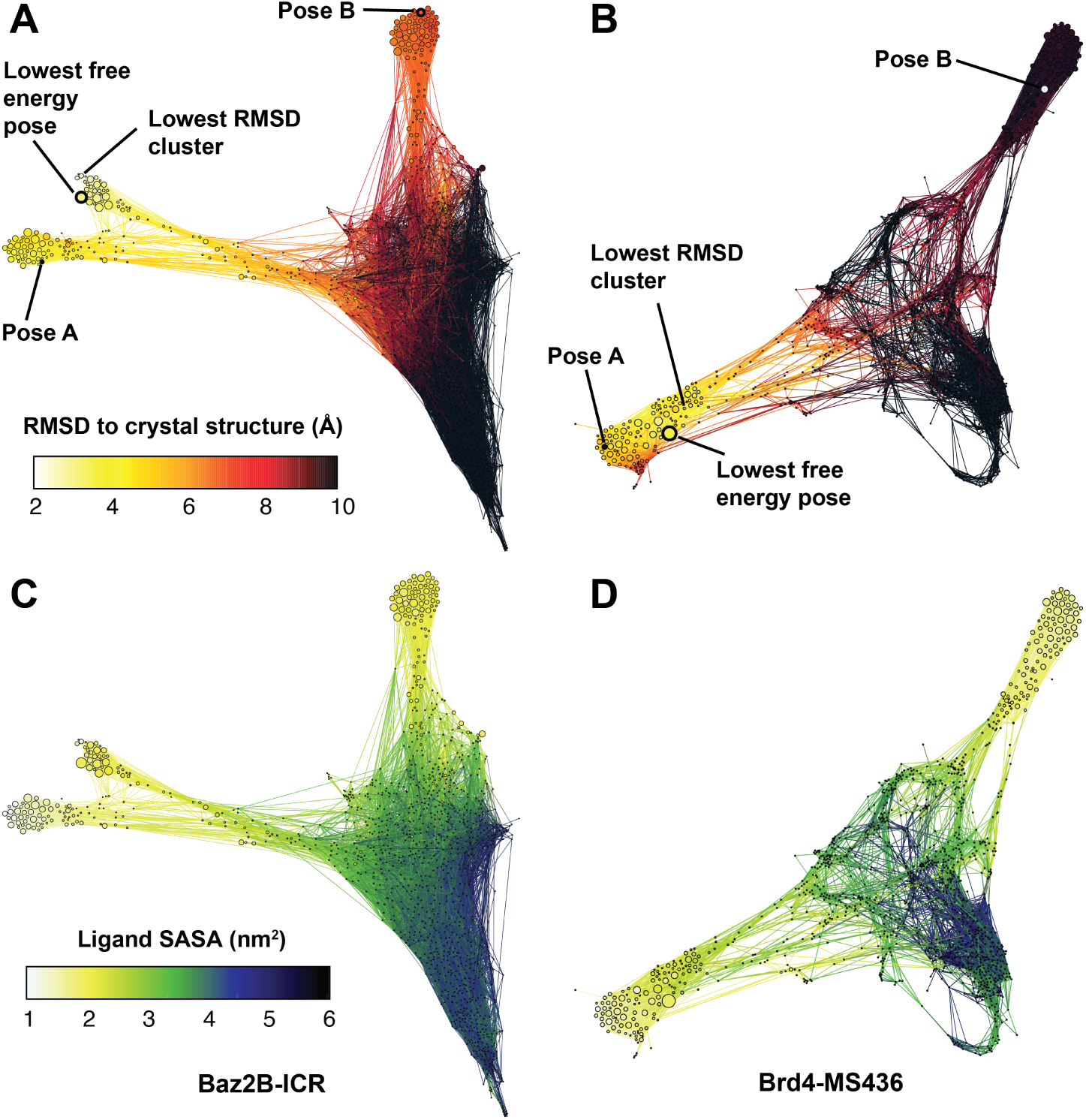
Networks of bound poses. The simulations starting from poses A and B are synthesized into networks where each node is a bound pose and the connections show the poses that are observed to interconvert in the molecular dynamics simulations. In all networks, nodes are given a size according to their weight, as determined by the Markov state model with *n_tICA_* = 5, *τ_tICA_* = 0.2 ns, *n_c_* = 1200, *p_excl_* = 70 and *τ* = 0.2 ns. (**A,B**) Nodes are colored according to their RMSD to the crystal structure, determined using the average over 10 randomly chosen structures from each node. The clusters corresponding to starting poses A and B are labeled for each network, as well as the predicted lowest free energy pose for this set of Markov state model parameters. (**C,D**) Nodes are colored according to their ligand solvent accessible surface area (SASA). In both networks, quasi-bound (green) and unbound (blue/black) nodes form a dense cluster, with deeply bound states (white/yellow) radiating outward from this cluster.

Figures 6A and 6B are colored according to the RMSD of the ligand to the crystal structure after aligning to the C*α* atoms in the binding site. In both cases, starting pose A has much lower RMSD to the crystal than pose B, as shown in Figure 1. The lowest free energy pose for this set of MSM parameters is indicated in both cases, as described in Section III C 3. Due to the nature of the network layout minimization algorithm, nodes that are far apart tend to interconvert more slowly than nodes that are close together. Tight groups of nodes can thus be considered to be in the same basin of attraction, and to interconvert relatively quickly. For both systems we find the lowest free-energy pose to be in the same basin of attraction as the lowest RMSD pose to the crystal structure.

Figures 6C and 6D are colored according to the ligand solvent accessible surface area (SASA). From these we can clearly identify three deeply bound basins of attraction for Baz2B-ICR7 (yellow and white nodes), and two deeply bound basins for Brd4-MS436. The “Pose A basin” for Baz2B-ICR7 corresponded to a deeper insertion of the ICR7 ligand, with roughly the same orientation as the crystal structure. The network shows that the crystal structure basin is “off-pathway” with respect to “Pose A” binding and unbinding transitions. The Brd4-MS436 “Pose A basin” is also more deeply bound than the crystal structure, and the lowest free energy pose. However, the crystal structure in this case is “on-pathway” with respect to “Pose A” binding and unbinding transitions.

From the SASA network plots it is also apparent how many distinct local minima exist with relatively similar predicted probabilities. The lowest free energy poses have probabilities of 0.015 and 0.027 for Baz2B and Brd4, respectively. For Baz2B there are 25 other states with at least half of this probability, with RMSDs to crystal ranging from 2.9 to 9.8 Å. For Brd4 there are 7 other states that are at least half as probable, with RMSDs to crystal ranging from 3.0 to 9.9 Å. This abundance of states with similar stabilities underscores the challenge of conclusively predicting a single stable pose.

### E. Pose interconversion pathways

The pose networks provide a detailed description not only of binding and unbinding pathways, but the interconversion of different states. When a ligand interconverts between different bound poses, will this occur while the ligand remains loosely associated with the protein? Or will the ligand first unbind, and then rebind in a different orientation? To answer this question we label a set of nodes as “unbound” if the minimum distance between the ligand and the protein is greater than 5 Å. Note that this must be smaller than the cutoff we use for terminating unbinding trajectories (10 Å). We then divide the remainder of each network into communities using a modularity optimization algorithm implemented in Gephi [54]. An extended transition matrix (*T_α_*) is constructed (size 2*N* x 2*N*, where *N* = 1200 is the number of states) using an auxiliary variable *α*, where *α* = 0 if a trajectory has not yet visited the unbound state, and *α* = 1 indicates a trajectory *has* visited the unbound state. To examine transition paths between communities *A* and *B*, we add probability sinks to *T_α_* by replacing columns that correspond to *A* and *B* with identity vectors (*i.e.* probability can get in, but cannot get out). By iteratively multiplying this matrix against itself we can examine its steady state behavior, and for each state *i*, measure the committor probabilities 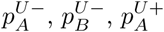, and 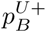, where the subscript indicates the destination basin, and the superscript indicates if the transition path did (*U* +) or did not (*U* −) go through the unbound state. The probability of an *A* to *B* transition path being mediated by the unbound state is then equal to:
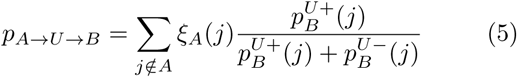

where *ξ_A_*(*i*) is the probability that a trajectory, initialized in basin *A*, will exit into state *i*. The exit probabilities (*ξ_A_*(*i*)) are calculated again using a sink matrix, this time using a sink for each state that is not in basin *A*. This analysis (referred to as “hub score analysis”) was previously used to study interconversion paths in protein folding conformation space networks [52, 55], and is implemented in the CSNAnalysis package [53].

Figure 7 shows the communities in each network (A and B), and the probabilities that transition paths between these communities are mediated by the unbound state (C and D). In both networks the unbound states are shown as black nodes. It is important to note that the unbound states are not expected to form a cohesive interconverting community, as the ligand can unbind from anywhere on the protein, and a wide variety of unbound states are observed (as shown in Figure 2). For Baz2B, there are three “deeply bound” communities: one for each of “pose A” (1) and “pose B” (0), and an additional community that includes the lowest free energy state (Community 2). Brd4 has two deeply bound communities, corresponding to poses A and B. The community indices are ordered according to their total weight, with community 0 having the highest total weight.

**FIG. 7.**
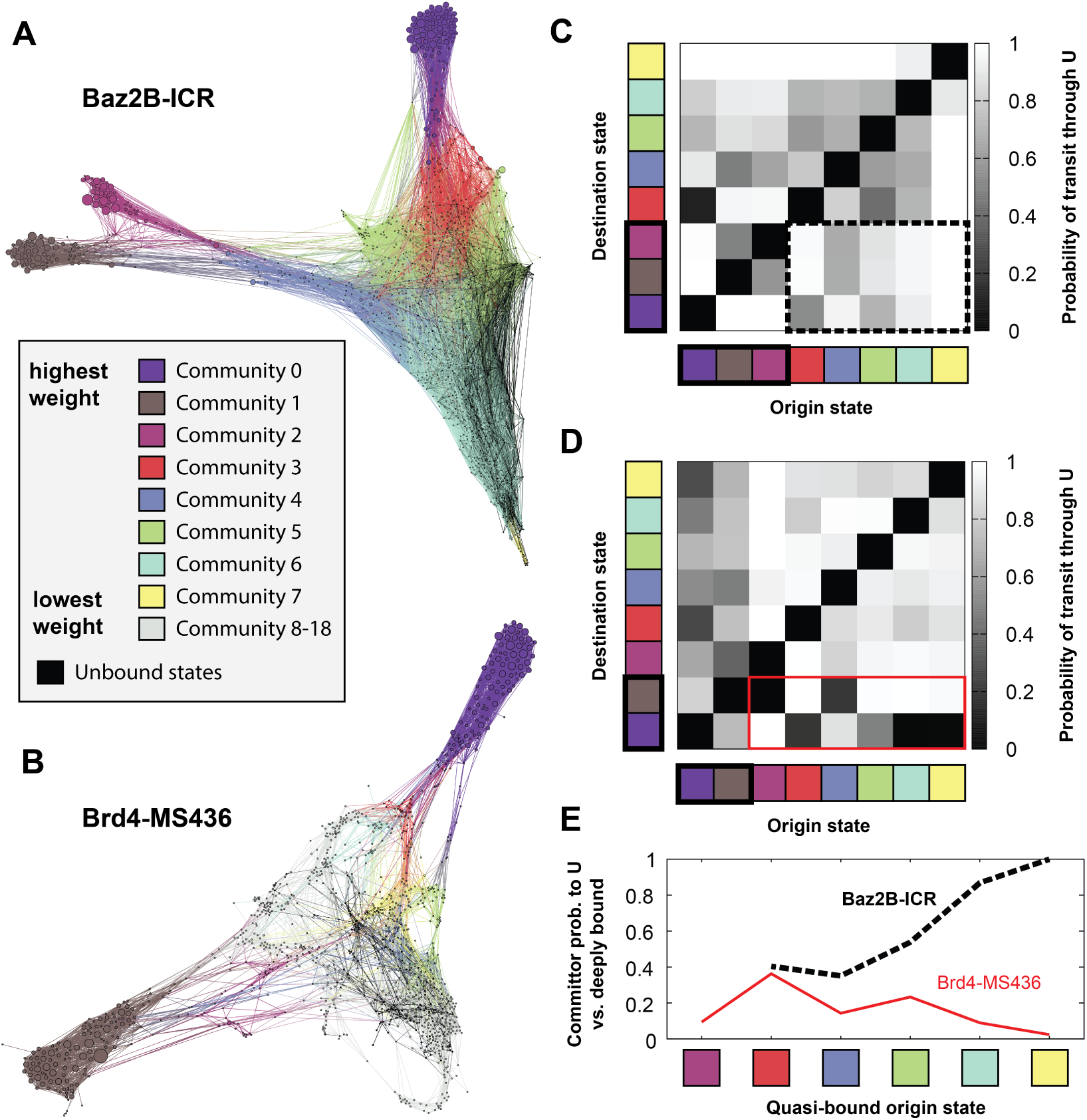
Bound-to-bound transition paths using the unbound state as a mediator. (**A,B**) Conformation space networks for Baz2B-ICR7 and Brd4-MS436, colored to show communities identified by modularity optimization. In both networks communities are ordered according to their total weight, with Community 0 (purple) having the highest weight. The unbound states are shown in black. In panel **B**, communities 8-18 are shown in grey and are not considered in further analysis. (**C,D**) Matrices of mediation probabilities are shown for both systems. Each element in the matrix shows the probability that a transition path from the origin state to the destination state uses the unbound state as a mediator. For Baz2B-ICR7 (**C**), the first three communities are labeled “deeply bound” and are highlighted with a black rectangle. The dotted line surrounds the mediation probabilities for “quasi-bound” to “deeply bound” transitions. For Brd4-MS436 (**D**), the first two communities are labeled “deeply bound”, and “quasi-bound” to “deeply bound” transitions are marked with a solid red line. Panel (**E**) compares the committor probability between unbound states and the set of deeply bound states for each quasi-bound starting state.

In the bottom left of the matrices in panels C and D we see the probabilities that interconversion paths between the highest weighted communities goes through the unbound state. For Baz2B interconversion between the “Pose B” community (“C0”, purple), with the “Pose A” community (“C1”, grey) almost always goes through the unbound state (*p* = 0.99). Similarly, paths from “Pose B” to C2 (magenta), which contains the crystal structure pose, go through the unbound state with *p* = 0.98. In the dashed rectangle, we see that paths from “quasi-bound” communities (C3-C7) to deeply bound communities go through the unbound state with high probability. Remarkably, even paths from C1 to C2 go through the unbound state with *p* = 0.504, and from C2 to C1 with *p* = 0.579. The lowest unbound mediation probability is for paths from C0 to C3 (*p* = 0.130), where C3 is directly between C0 and the unbound states. With this exception, we conclude that in the Baz2B network interconversion between poses mostly proceeds through unbinding and rebinding. Significant differences are observed for the Brd4 mediation probabilities. Each quasi-bound state has a deeply-bound state that it can transition to directly, without passing through the unbound states (Panel D: dark grey and black squares in the red rectangle). However, transitions between the two deeply-bound states are still mediated through unbound states more often than not (*p* = 0.820 for C0-C1 and *p* = 0.728 for C1-C0).

To further investigate the binding landscape we determine the probability that quasi-bound trajectories will commit to either one of the deeply-bound states, or to the unbound states. A committor probability is determined for each quasi-bound state, and a weighted average (using the equilibrium weights of each state) is shown in Figure 7E for each community. We see significant differences between the two bromodomain-inhibitor systems, with most of the Baz2 quasi-bound communities committing to the unbound state, and all of the Brd4 quasi-bound communities committing to deeply-bound states. The Brd4-MS436 system can thus be thought of as a “dual-funnel”-shaped landscape, with two minima near poses *A* and *B*. The quasi-bound states exist halfway down this funnel, and help guide the ligand to one of the two deeply bound orientations. In the Baz2B-ICR7 system the quasi-bound states have still not crossed the rate-limiting step of binding, and the ligand is still free to reorient, or not to bind at all.

## IV. DISCUSSION

Mapping the ligand binding landscape can teach us how ligands bind, how poses are connected, and which poses are the most stable. Here we have shown that WExplore is able to efficiently build a model of the binding landscape, and sample events with waiting times up to 280 ms (the release of MS436 from pose A). However, it is limited by the accuracy of the force field used for both the ligand and the protein. In particular, we suspect that the parameterization of ICR under-predicted the stability of the ring-stacked conformation. This could have led to the over-prediction of the *K_D_* in Table III, and the extremely short mean first passage times of ligand release (420 and 150 ns for poses A and B, respectively). In contrast, a Baz2B probe with 60 nM affinity, GSK2801, was measured using biolayer interferometry to have a *k*_off_ of 6.95 × 10^−3^ s^−1^, which would have a mean first passage time of 144 s [35].

Pose prediction by integrating unbinding simulations into a network model presents a novel approach to a difficult problem. Previously it was shown that long, straightforward simulations can discover binding poses. Shan et al. directly simulated binding of two inhibitors to Src kinase, obtaining 4 binding events in 150 *µ*s of total simulation [56]. Generally, ligand binding occurs more quickly than unbinding, and is thus more amenable to direct simulation. However the residence time of each pose, which can be obtained only through simulating ligand release, is also necessary to rank pose stability. Clark et al. used induced fit docking in combination with metadynamics to determine pose stability [57]. Ten trajectories were run for each pose, each 10 ns in length. The authors showed an improvement over induced fit docking alone, and the cost of this approach (100 ns per ligand per pose) is much cheaper than the approach presented here (2.1 *µ*s per pose). However, the metadynamics approach can only rank poses, and cannot discover new poses that were not in the induced docking set. It also requires the definition of an order parameter that is appropriate to use for each pose and each system, which is not straightforward. It is worth noting also that our combined network model approach becomes more accurate as more starting poses are added to the system. One could imagine, for example, adding short simulations from a large number of additional starting poses to the networks constructed here.

The finding that most poses interconvert through the unbound state has direct implications for the discovery of new bromodomain inhibitors. A common approach in virtual screening is to combine docking with MD simulations to evaluate binding pose stability [58-60]. This is typically done as a filter late in a screening pipeline, for instance to select the top 24 compounds in a set of 55 [61]. To assess the stability of binding poses, it is common to use trajectories that are tens to hundreds of nanoseconds in length. In the regime where poses interconvert directly without unbinding, a single stability measurement could report on multiple poses. However, as we find that different poses are typically connected through the unbound state, we would recommend to test multiple putative bound poses for each compound during screening, and discard only compounds that show no stable poses.

This finding has similar implications for kinetics-based drug design. In the regime where poses interconvert directly, all poses will have similar off-rates. Interconverting through the unbound state implies that a weighted average of pose-specific off-rates 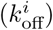 values should be used to determine an apparent off-rate 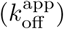:
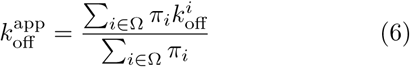

where the sum is over the set of all bound states (Ω) and *π_i_* is the probability of state *i*. If one bound state is much more populated than the others (e.g. *π_i_* ≫ *π_j_* ∀ *j* ≠ *i*) then 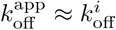.

Mapping the binding landscape could also aid in our understanding of biological processes. By studying the landscape of natural protein-ligand association processes, such as enzyme substrates, we can observe whether proteins have evolved to maximize the fraction of successful binding encounters. Nature-inspired strategies to maximize binding rates could be useful to help improve binding affinities, as well as improve the selectivity profiles of covalent inhibitors.

## V. ACKNOWLEDGEMENTS

ARD would like to thank Samuel D. Lotz for contributions to the wepy package [51] that enabled the variable lag time analysis used here. This work was supported by NSF grant DMS1761320.

## References

[1] R. A. Friesner, J. L. Banks, R. B. Murphy, T. A. Halgren, J. J. Klicic, D. T. Mainz, M. P. Repasky, E. H. Knoll, M. Shelley, J. K. Perry, D. E. Shaw, P. Francis, and P. S. Shenkin, Journal of Medicinal Chemistry 47, 1739 (2004).

[2] H. Chen, P. D. Lyne, F. Giordanetto, T. Lovell, and J. Li, Journal of Chemical Information and Modeling 46, 401 (2006).

[3] O. Trott and A. Olson, Journal of Computational Chemistry 31, 455 (2010).

[4] D. Fox, A. B. Burgin, and M. E. Gurney, Cellular Signalling 26, 657 (2014).

[5] H. Lu, K. England, C. A. Ende, J. J. Truglio, S. Luckner, B. G. Reddy, N. L. Marlenee, S. E. Knudson, D. L. Knudson, R. A. Bowen, C. Kisker, R. A. Slayden, and P. J. Tonge, ACS Chemical Biology 4, 221 (2009).

[6] S. R. Luckner, N. Liu, C. W. Am Ende, P. J. Tonge, and C. Kisker, Journal of Biological Chemistry 285, 14330 (2010).

[7] D. Guo, L. H. Heitman, and A. P. Ijzerman, ACS Medicinal Chemistry Letters 7, 819 (2016).

[8] D. A. Schuetz, W. E. A. de Witte, Y. C. Wong, B. Knasmueller, L. Richter, D. B. Kokh, S. K. Sadiq, R. Bosma, I. Nederpelt, L. H. Heitman, E. Segala, M. Amaral, D. Guo, D. Andres, V. Georgi, L. A. Stoddart, S. Hill, R. M. Cooke, C. De Graaf, R. Leurs, M. Frech, R. C. Wade, E. C. M. de Lange, A. P. IJzerman, A. Müller-Fahrnow, and G. F. Ecker, Drug Discovery Today 22, 896 (2017).

[9] D. Huang and A. Caflisch, PLoS Computational Biology 7, e1002002 (2011).

[10] A. Dickson, P. Tiwary, and H. Vashisth, Current Topics in Medicinal Chemistry 17 (2017).

[11] D. B. Kokh, M. Amaral, J. Bomke, U. Grädler, D. Musil, H.-P. Buchstaller, M. K. Dreyer, M. Frech, M. Lowinski, F. Vallée, M. Bianciotto, A. Rak, and R. C. Wade, Journal of Chemical Theory and Computation (2018).

[12] A. Laio and M. Parrinello, Proc. Nat. Acad. Sci. USA 99, 12562 (2002).

[13] P. Tiwary, J. Mondal, and B. J. Berne, Science Advances 3, e1700014 (2017).

[14] R. Casasnovas, V. Limongelli, P. Tiwary, P. Carloni, and M. Parrinello, Journal of the American Chemical Society 139, 4780 (2017).

[15] P. Tiwary and M. Parrinello, Physical Review Letters 111, 230602 (2013).

[16] F. Cérou and A. Guyader, Stochastic Analysis and Applications 25, 417 (2007).

[17] I. Teo, C. G. Mayne, K. Schulten, and T. Lelièvre, Journal of Chemical Theory and Computation 12, 2983 (2016).

[18] S. Doerr and G. De Fabritiis, Journal of Chemical Theory and Computation 10, 2064 (2014).

[19] G. G. A. Huber and S. Kim, Biophysical Journal 70, 97 (1996).

[20] S. H. Northrup, S. A. Allison, and J. A. McCammon, The Journal of Chemical Physics 80, 1517 (1984).

[21] A. Rojnuckarin, D. R. Livesay, and S. Subramaniam, Biophysical Journal 79, 686 (2000).

[22] A. S. Saglam and L. T. Chong, The Journal of Physical Chemistry B 120, 117 (2015).

[23] A. Dickson and C. L. Brooks III, The Journal of Physical Chemistry B 118, 3532 (2014).

[24] A. Dickson and S. D. Lotz, Journal of Physical Chemistry B 120, 5377 (2016).

[25] A. Dickson and S. D. Lotz, Biophysical Journal 112, 620 (2017).

[26] T. Dixon, A. Dickson, and S. D. Lotz, Submitted (2018).

[27] P. Filippakopoulos and S. Knapp, Nature Reviews Drug Discovery 13, 337 (2014).

[28] F. M. Ferguson, O. Fedorov, A. Chaikuad, M. Philpott, J. R. C. Muniz, I. Felletar, F. Von Delft, T. Heightman, S. Knapp, C. Abell, and A. Ciulli, Journal of Medicinal Chemistry 56, 10183 (2013).

[29] G. Zhang, A. N. Plotnikov, E. Rusinova, T. Shen, K. Morohashi, J. Joshua, L. Zeng, S. Mujtaba, M. Ohlmeyer, and M. M. Zhou, Journal of Medicinal Chemistry 56, 9251 (2013).

[30] A. Chaidos, V. Caputo, and A. Karadimitris, Therapeutic Advances in Hematology 6, 128 (2015).

[31] J. P. Daguer, C. Zambaldo, D. Abegg, S. Barluenga, C. Tallant, S. Müller, A. Adibekian, and N. Winssinger, Angewandte Chemie - International Edition 54, 6057 (2015).

[32] K. Dhananjayan, Journal of Cancer Research 2015, 1 (2015).

[33] L. Drouin, S. McGrath, L. R. Vidler, A. Chaikuad, O. Monteiro, C. Tallant, M. Philpott, C. Rogers, O. Fedorov, M. Liu, W. Akhtar, A. Hayes, F. Raynaud, S. Müller, S. Knapp, and S. Hoelder, Journal of Medicinal Chemistry 58, 2553 (2015).

[34] A. K. Urick, L. M. Hawk, M. K. Cassel, N. K. Mishra, S. Liu, N. Adhikari, W. Zhang, C. O. Dos Santos, J. L. Hall, and W. C. Pomerantz, ACS Chemical Biology 10, 2246 (2015).

[35] P. Chen, A. Chaikuad, P. Bamborough, M. Bantscheff, C. Bountra, C. W. Chung, O. Fedorov, P. Grandi, D. Jung, R. Lesniak, M. Lindon, S. Müller, M. Philpott, R. Prinjha, C. Rogers, C. Selenski, C. Tallant, T. Werner, T. M. Willson, S. Knapp, and D. H. Drewry, Journal of Medicinal Chemistry 59, 1410 (2016).

[36] T. D. Crawford, V. Tsui, E. M. Flynn, S. Wang, A. M. Taylor, A. Cote, J. E. Audia, M. H. Beresini, D. J. Burdick, R. Cummings, L. A. Dakin, M. Duplessis, A. C. Good, M. C. Hewitt, H. R. Huang, H. Jayaram, J. R. Kiefer, Y. Jiang, J. Murray, C. G. Nasveschuk, E. Pardo, F. Poy, F. A. Romero, Y. Tang, J. Wang, Z. Xu, L. E. Zawadzke, X. Zhu, B. K. Albrecht, S. R. Magnuson, S. Bellon, and A. G. Cochran, Journal of Medicinal Chemistry 59, 5391 (2016).

[37] J.-R. R. Marchand, G. Lolli, and A. Caflisch, Journal of Medicinal Chemistry 59, 9919 (2016).

[38] U. Raj, H. Kumar, and P. Varadwaj, Journal of Biomolecular Structure and Dynamics 1102, 1 (2016).

[39] R. P. Nowak, S. L. Deangelo, D. Buckley, Z. He, K. A. Donovan, J. An, N. Safaee, M. P. Jedrychowski, C. M. Ponthier, M. Ishoey, T. Zhang, J. D. Mancias, N. S. Gray, J. E. Bradner, and E. S. Fischer, Nature Chemical Biology (2018).

[40] P. Filippakopoulos, J. Qi, S. Picaud, Y. Shen, W. B. Smith, O. Fedorov, E. M. Morse, T. Keates, T. T. Hickman, I. Felletar, M. Philpott, S. Munro, M. R. McKeown, Y. Wang, A. L. Christie, N. West, M. J. Cameron, B. Schwartz, T. D. Heightman, N. La Thangue, C. A. French, O. Wiest, A. L. Kung, S. Knapp, and J. E. Bradner, Nature 468, 1067 (2010).

[41] K. Vanommeslaeghe and A. D. MacKerell, Journal of Chemical Information and Modeling 52, 3144 (2012).

[42] K. Vanommeslaeghe, E. Prabhu Raman, and A. D. MacKerell, Journal of Chemical Information and Modeling 52, 3155 (2012).

[43] B. W. Zhang, D. Jasnow, and D. M. Zuckerman, Journal of Chemical Physics 132, 1 (2010).

[44] M. C. Zwier, J. W. Kaus, and L. T. Chong, Journal of Chemical Theory and Computation 7, 1189 (2011).

[45] E. N. Laricheva, G. B. Goh, A. Dickson, and C. L. Brooks III, Journal of the American Chemical Society 137, 2892 (2015).

[46] A. Dickson, A. Mustoe, L. Salmon, and C. Brooks III, Nucleic Acids Research 42, 12126 (2014).

[47] S. D. Lotz and A. Dickson, Journal of the American Chemical Society 140, 618 (2018).

[48] A. Dickson, A. Warmflash, and A. R. Dinner, Journal of Chemical Physics 131, 154104 (2009).

[49] C. R. Schwantes and V. S. Pande, Journal of Chemical Theory and Computation 9, 2000 (2013).

[50] G. Pérez-Hernández, F. Paul, T. Giorgino, G. De Fabritiis, and F. Noé, Journal of Chemical Physics 139 (2013).

[51] A. Dickson, S. Lotz, A. Uyar, N. Donyapour, and T. Dixon, https://github.com/ADicksonLab/wepy “wepy,” (2018).

[52] A. Dickson and C. L. Brooks III, Journal of the American Chemical Society 135, 4729 (2013).

[53] A. Dickson, https://github.com/ADicksonLab/CSNAnalysis “CSNAnalysis,” (2018).

[54] M. Bastian, S. Heymann, and M. Jacomy (AAAI Press, 2009).

[55] A. Dickson and C. L. Brooks III, Journal of Chemical Theory and Computation 8, 3044 (2012).

[56] Y. Shan, E. T. Kim, M. P. Eastwood, R. O. Dror, M. A. Seeliger, and D. E. Shaw, Journal of the American Chemical Society 133, 9181 (2011).

[57] A. J. Clark, P. Tiwary, K. Borrelli, S. Feng, E. B. Miller, R. Abel, R. A. Friesner, and B. J. Berne, Journal of Chemical Theory and Computation 12, 2990 (2016).

[58] A. Cavalli, G. Bottegoni, C. Raco, M. De Vivo, and M. Recanatini, Journal of Medicinal Chemistry 47, 3991 (2004).

[59] T. Sakano, M. Mahamood, T. Yamashita, and H. Fuijitani, Biophysics and Physicobiology 13, 181 (2016).

[60] K. Liu, E. Watanabe, and H. Kokubo, Journal of Computer-Aided Molecular Design 31, 201 (2017).

[61] H. Zhao and A. Caflisch, European Journal of Medicinal Chemistry 91, 4 (2014).

